# CRISPR/Cas9-mediated gene disruption of endogenous co-receptors confers broad resistance to HIV-1 in human primary cells and humanized mice

**DOI:** 10.1101/2021.06.30.450601

**Authors:** Shasha Li, Leo Holguin, John C. Burnett

## Abstract

In this project, we investigated the CRISPR/Cas9 system for creating HIV resistance by targeting the human *CCR5* and *CXCR4* genes, which encode cellular co-receptors required for HIV-1 infection. Using a clinically scalable system for transient *ex vivo* delivery of Cas9/gRNA ribonucleoprotein (RNP) complexes, we demonstrated that CRISPR-mediated disruption of *CCR5* and *CXCR4* in T-lymphocytes cells significantly reduced surface expression of the co-receptors, thereby establishing resistance to HIV-1 infection by CCR5 (R5)-tropic, CXCR4 (X4)-tropic, and dual (R5/X4)-tropic strains. CRISPR-mediated disruption of the *CCR5* alleles in human CD34^+^ hematopoietic stem and progenitor cells (HSPCs) led to the differentiation of HIV-resistant macrophages. In human CD4^+^ T cells transplanted into a humanized mouse model, disruption of *CXCR4* inhibited replication of X4-tropic HIV-1, thus leading to the virus-mediated enrichment *CXCR4*-disrupted cells in the peripheral blood and spleen. However, in human CD4^+^ T cells with both *CCR5* and *CXCR4* disruption, we observed poor engraftment in bone marrow, although significant changes were not observed in the lung, spleen, or peripheral blood. This study establishes a clinically scalable strategy for the dual knockout of HIV-1 co-receptors as a therapeutic strategy, while also raising caution of disrupting *CXCR4*, which may abate engraftment of CD4^+^ T cells in bone marrow.

## Introduction

Human immunodeficiency virus 1 (HIV-1), the virus that causes AIDS, currently afflicts more than 38 million people worldwide.^1^ Despite the effectiveness of antiretroviral therapy (ART) in controlling HIV-1 replication and infection, these drugs are unable to eradicate the virus from a patient. Complicating matters, accessibility to ART and daily compliance are challenging for millions living with HIV, and HIV-infected individuals disproportionately suffer from accelerated aging and an increased risk of age-related health complications.^2^ Unfortunately, An HIV-1 cure remains elusive. Innovative therapeutic strategies are currently being explored as potential alternatives to ART,^3^ including gene-editing strategies which inhibit viral infection.^4^

The HIV-1 replication cycle begins with the viral particle binding to the CD4 receptor and then either to the CCR5 or the CXCR4 co-receptor on the target cells. Binding then triggers fusion of the viral and host cell membranes, thereby facilitating entry into the cell, where the viral genome undergoes reverse transcription and integration into the host genome. Of the two primary co-receptors, CCR5 is the cellular co-receptor used by the majority of HIV-1 strains for binding and entry ^5^ and is critical for primary infection via mucosal transmission.^6^ Approximately ∼1% of individuals of northern European descent are homozygous for the *CCR5Δ32* allele, which is characterized by a 32-bp deletion that results in a truncated CCR5 protein that is not expressed on the cell surface. While these individuals are healthy despite lacking a functional *CCR5* gene, they are also highly resistant to HIV-1 infection.^7,8^ The first two documented functional cures of HIV-1 were with patients who received allogeneic transplantation with hematopoietic stem cells from *CCR5Δ32* homozygous donors for the treatment of acute myeloid leukemia^9,10^ or refractory Hodgkin lymphoma.^11^ However, this general strategy has been met with mixed success, and several other patients have experienced complications due to allogeneic stem cell transplantation or relapse of underlying cancer,^12,13^ while others have been marked by the emergence of CXCR4 (X4)-tropic HIV-1 strains that do not utilize the CCR5 co-receptor.^14^

Numerous gene editing tools have been used against *CCR5* to inhibit R5-tropic HIV-1 infection *in vitro* and *in vivo*, including ZFN,^15-18^ TALEN,^19-21^ and CRISPR/Cas systems.^22-24^ Due to the possibility of HIV-resistance to CCR5 gene disruption, which occurs through natural tropism shift, it is likely necessary to disrupt CXCR4 to eradicate HIV-1 infections in most individuals. Hence, ZFN^25,26^ and CRISPR/Cas^27,28^ systems have been designed edit *CXCR4* for the inhibition of X4-tropic HIV-1. Moreover, a few studies have explored the simultaneous disruption of CCR5 and CXCR4 alleles using two zinc-finger nucleases^29^ or two sgRNAs via CRISPR/Cas9.^30^ Although many of these approaches are still in the preclinical stage, clinical trials primarily focused on the use of ZFN^31,32^ or CRISPR/Cas9^33^ for *CCR5* editing have yielded promising results in clinical safety and efficacy tests, while *CXCR4* gene editing strategies have not yet been tested clinically.

Translation of gene editing technology utilizing disrupting co-receptors for treating HIV/AIDS, demands exquisite on-target precision, ample efficiency, and delivery approaches that are scalable and clinically feasible. In the present study, we have utilized the CRISPR-Cas9 gene-editing system to disrupt *CCR5, CXCR4* genes or both to create HIV-resistance in human primary T cells in a clinically scalable system. Importantly, we demonstrated that the resulting cells yield different selective advantages in HIV infection, with specific HIV-1 strains (R5 tropic, X4 tropic and dual tropic) that utilize either the *CCR5* or *CXCR4* surface receptors or both. Next, we evaluated the gene-modified cells in a humanized mouse model. Our study gives an in-depth investigation of CRISPR disrupted CCR5 or/and CXCR4 co-receptor in aim of a cure for HIV/AIDS. These experiments lay the groundwork for creating HIV-1 resistance in a clinically scalable system.

## Results

### CRISPR-Cas9-mediated disruption of CCR5 protect cells from HIV-1 infection

To evaluate the CRISPR-Cas9 system in creating HIV-resistant cells, we first utilized a previously described approach with lentiviral expression of both the single guide RNA (sgRNA) and human codon-optimized *Streptococcus pyogenes* Cas9 (spCas9) components, as well as the TagRFP reporter gene.^34^ Using a sgRNA design algorithm,^35^ we selected unique guides sequences to target *CCR5* with the CRISPR-Cas9 system. CEM.NKRCCR5+ cells (i.e., human CD4^+^ lymphoblast cells with retroviral vector expression of human CCR5^36^) were transduced with the lentiviral vectors at a low multiplicity of infection (MOI ∼ 0.1). A control vector was created which carried an irrelevant sgRNA sequence in addition to the spCas9 and TagRFP expression cassettes. One week after transduction, transduced cells were sorted by FACS for TagRFP expression and analyzed for CCR5 surface expression by flow cytometry to assess the degree of CRISPR-mediated gene knockout. Surface expression of CCR5 was significantly reduced in the cells treated with CCR5-CRISPR (81.7% CCR5^+^ cells in control vs. 4.3% CCR5^+^ cells in CCR5-CRISPR, **Figure 1A**). Genomic DNA was analyzed for gene editing using CEL1 Surveyor Nuclease Assay, which revealed 62% ablation efficiency of *CCR5* (**Figure 1B**). Gene disruption was further characterized by NGS analysis of across the *CCR5* target site, which revealed significant and frequent insertions and deletions (indels) at the sgRNA target site, consistent with the imprecise DNA repair mechanism of non-homologous end-joining (NHEJ) (**Figure S1)**. Deep sequencing of the *CCR5* target site revealed CRISPR-induced indels in 87.9% of the total reads (**Figure 1C**), ranging from single base pair (bp) insertions or deletions to insertions or deletions exceeding 100bp. To investigate whether CRISPR-mediated disruption of the *CCR5* gene facilitated HIV resistance, we challenged the gene-modified CEM cells with R5-tropic HIV-1_Bal_ and observed HIV replication over a 4-week time course. HIV-1 replication was suppressed in the CCR5-CRISPR cells, with supernatant p24 antigen levels greater than 100-fold lower than the control group at 14, 17, 21, and 28 days after HIV-1_BaL_ challenge (**Figure 1D**).

**Figure 1.**
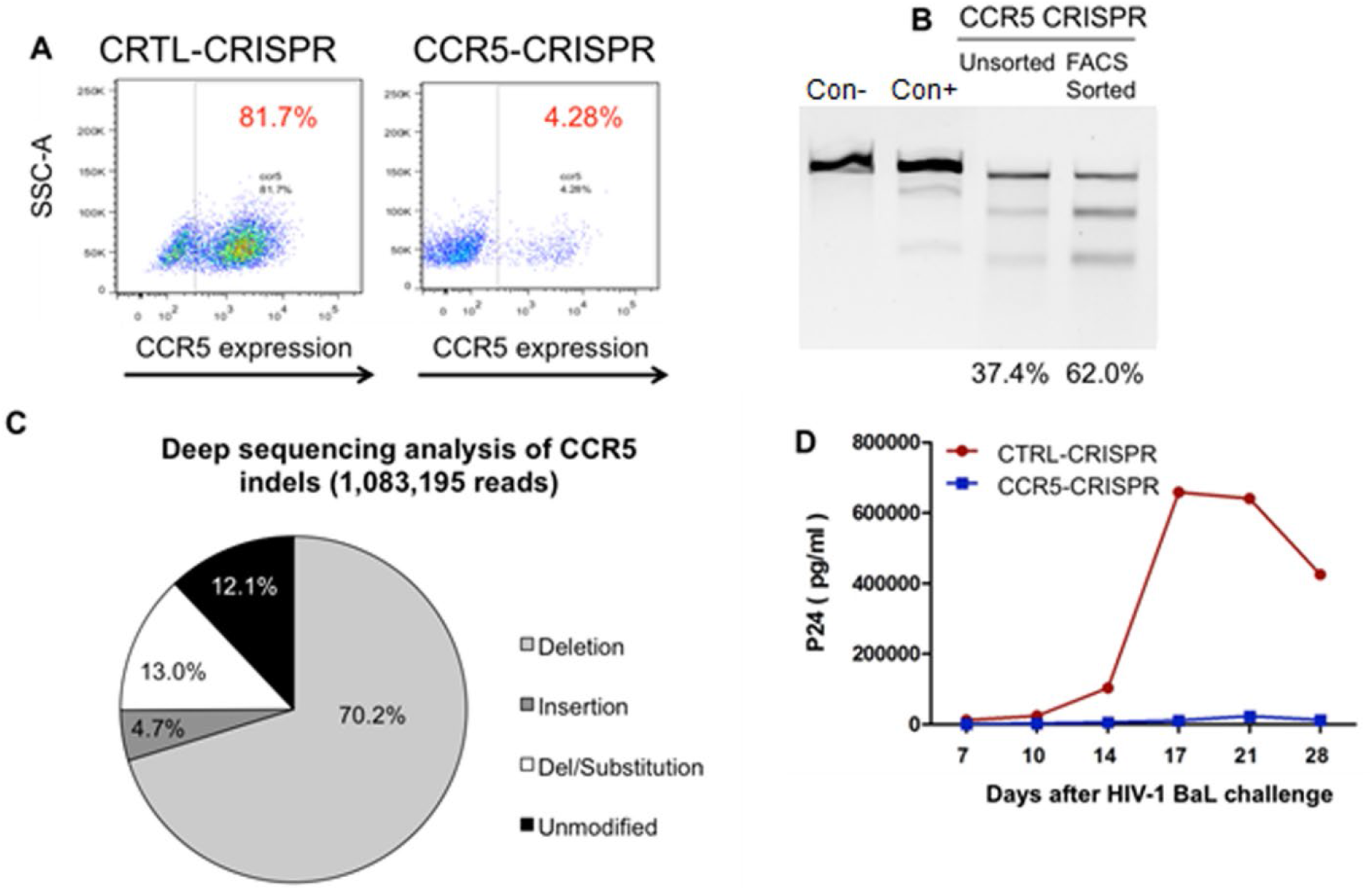
CRISPR/Cas9 gene disruption of CCR5 in CD4^+^ T cells. **A**. CEM CCR5^+^CD4^+^ T cells were transduced with CCR5-CRISPR vector or control vector and analyzed for CCR5 surface expression by flow cytometry. **B**. Indels detection by Surveyor assay in CEM CCR5^+^CD4^+^ T cells after CCR5 CRISPR-Cas9 modification. **C**. Deep sequencing analysis of CRISPR/Cas9-induced genome disruption by of CCR5. **D**. HIV-1_BaL_ replication in CEM CCR5+ CD4^+^ T cells treated with CCR5-CRISPR, as measured by HIV-1 p24 antigen in supernatant.

### HIV-1 resistance of CCR5 CRISPR-Cas9-modified CD34^+^ differentiated macrophages to CCR5 tropic HIV-1

R5-tropic HIV-1 strains (e.g., HIV-1_BaL_) are historically referred to as macrophage-tropic (M-tropic), as they are capable of infecting macrophages by utilizing the CCR5 co-receptor in addition to the CD4 receptor. Thus, we evaluated the antiviral efficacy of CRISPR-mediated gene disruption of CCR5 in primary macrophages that were derived from CD34^+^ hematopoietic stem and progenitor (HSPC) cells. Human CD34^+^ HSPCs were isolated from cord blood, transduced with the CCR5-CRISPR or control CRISPR lentiviral vectors, and sorted by FACS based on TagRFP expression (**Figure 2A**). The TagRFP-expressing CD34^+^ cells were differentiated into macrophages, as described in the Methods and Materials section (**Figure 2B)**. Macrophages were then challenged with HIV-1_BaL_ and evaluated for viral replication by p24 ELISA measurement of the supernatants over 28 days. HIV-1 replication was suppressed at all time points in macrophages treated with CCR5-CRISPR relative to control, with p24 antigen levels reduced greater than 10-fold at days 3, 7, and 21, greater than 25-fold in viremia at day 14, and greater than 5-fold at day 28 (**Figure 2C**). These results demonstrate that CRISPR-mediated disruption of *CCR5* in CD34^+^ HSPC-derived macrophages confers resistance to HIV-1 infection and replication.

**Figure 2.**
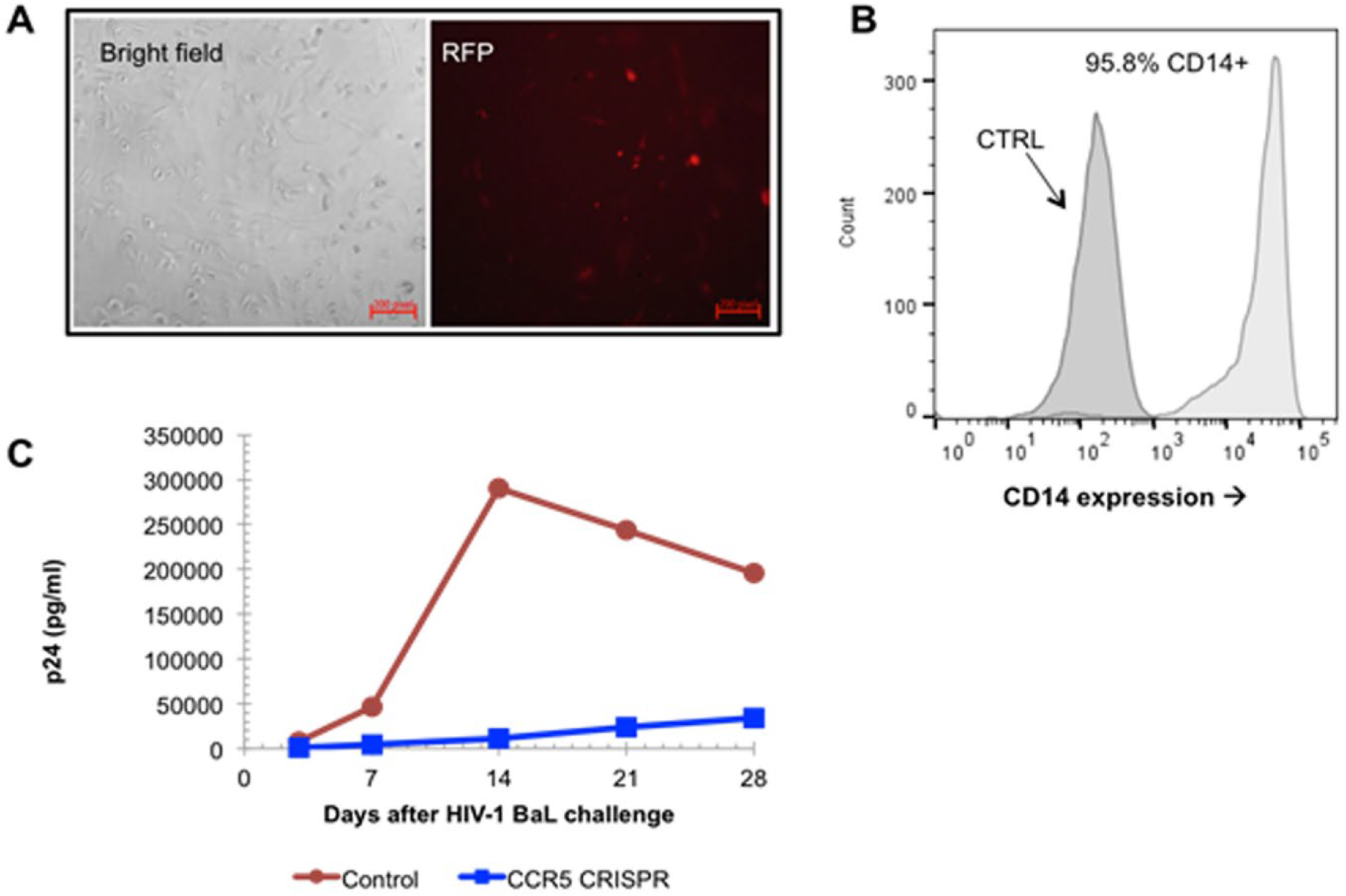
Hematopoietic differentiation and HIV resistance of macrophages from CRISPR-Cas9-CCR5 modified HSPCs. **A**. Morphology of macrophages generated from parental and CRISPR modified HSPCs. RFP indicated cells transduced with the CCR5-CRISPR vectors in differentiated macrophages. **B**. Flow cytometric analysis of macrophage-specific markers in macrophages generated from CRISPR modified HSPCs. **C**. Resistance of macrophage from CCR5-CRISPR modified HSPCs to HIV-1 virus infection comparing to unmodified cells.

### CRISPR-Cas9 gene disruption of CXCR4 confers resistance to T-tropic HIV-1 in cell lines and primary T cells

Although CRISPR-mediated disruption of the *CCR5* gene may confer resistance to R5-tropic HIV-1, it may not inhibit strains that utilize the CXCR4 (X4-tropic) or both CXCR4 and CCR5 co-receptors (dual-tropic). Thus, we designed guide CRISPR RNA sequences targeting *CXCR4* as an approach for inhibiting X4-tropic HIV-1. We first compared the efficacy of different sgRNA for each target, delivered using lentiviral vectors to disrupt surface CXCR4 expression on Jurkat CD4^+^ T cells. Flow cytometry analysis revealed a significant decrease in surface CXCR4 expression, with 99.3% CXCR4^+^ cells in control-CRISPR cells only 15.4% CXCR4^+^ cells transduced with CXCR4-CRISPR (**Figure 3A**). These observations were corroborated with analysis of editing of genome DNA by Surveyor nuclease assay, with 30.4% allelic disruption after CXCR4-CRISPR transduction (**Figure 3B**). Next, we assessed the biological effects of *CXCR4* disruption on preventing replication of X4-tropic HIV-1 in human PBMCs. Over a 16-day time course following HIV-1_NL4-3_ challenge, we observed significant resistance (p<0.05) to HIV replication in the CXCR4-CRISPR cells as measured by ELISA of supernatant at the indicated time points (**Figure 3C**). Collectively, these experiments demonstrate the feasibility of using CRISPR/Cas9 to engineer HIV resistant cells by targeting the *CCR5* and *CXCR4* host receptor genes.

**Figure 3.**
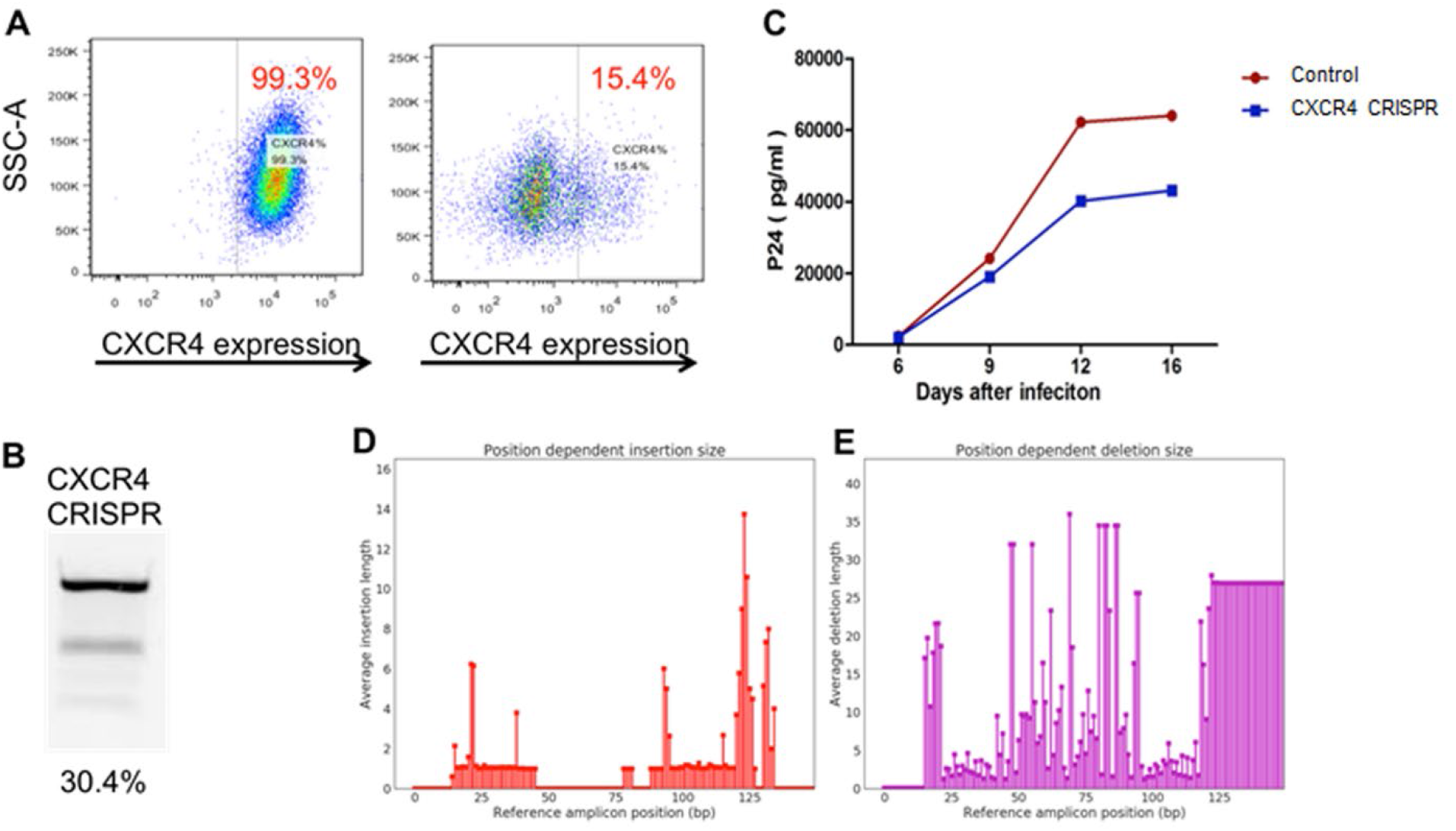
CRISPR/Cas9 gene disruption of CXCR4 in Jurkat T cells. **A**. Jurkat cells were transduced with CXCR4 CRISPR or control vector and analyzed for CXCR4 surface expression by flow cytometry. **B**. Surveyor nuclease assay detects indels in CXCR4 CRISPR modified Jurkat cells. **C**. HIV-1_NL4-3_ replication in PBMC treated with CXCR4 CRISPR, as measured by p24 in supernatant. **D**. Insertions or E. Deletions within the CXCR4 target site, as detected by Sanger sequencing and analyzed by inference of CRISPR edits (ICE). Insertions range from 1-14 bp while deletions range from 1-36 bp, as indicated by the y-axes.

### CXCR4 CRISPR-edited primary CD4^+^ T cells are selected in Hu-PBMC mice after infection with HIV-1 virus

While lentiviral delivery of the CRISPR-Cas9 system can achieve on-target efficacy, constitutive expression of the Cas9 and sgRNA components is also associated with high frequencies of off-target editing and is thus not suitable for clinical applications^37^. As an alternative delivery system, recombinant Cas9 protein may be complexed with the guide RNA for *ex vivo* delivery into cells by transient transfection or electroporation. The Cas9/gRNA ribonucleoprotein (RNP) provides burst-like kinetics that maximize the on-target efficiency, while minimizing less kinetically favorable off-target events^38^. Thus, we elected to deliver the Cas9 RNP to human primary CD4^+^ T cells using MaxCyte STX electroporation (MaxCyte, Inc.), as a similar approach has been previously demonstrated for the preparation of zinc finger nuclease-mediated gene-edited T cells at a clinical scale^39^. Specifically, we utilized the Alt-R CRISPR-Cas9 system (Integrated DNA Technologies, Inc.), which consists of spCas9 recombinant protein complexed with a trans-activating crRNA (tracrRNA) and a chemically modified CRISPR RNA (crRNA) that is specific for *CXCR4*. We utilized the human peripheral blood mononuclear cell (hu-PBMC) NSG mouse model to evaluate whether knockout of *CXCR4* in CD4^+^ T cells could protect cells *in vivo* from infection with X4-tropic HIV-1_NL4-3_ (**Figure 4A**). Two days after MaxCyte electroporation of AltR-CXCR4 CRISPR into human primary CD4^+^ T cells, flow cytometry analysis revealed that the subpopulation of CXCR4-negative T cells had increased from 2.3% to 20.2% in the CXCR4-CRISPR group (**Figure 4B**). Editing of the *CXCR4* alleles was also confirmed by Surveyor assay, which revealed 46% gene disruption (**Figure 4C**). Mice were analyzed for engraftment at 14 days after transplantation and were challenged with HIV-1_NL4-3_ at 28 days after transplantation. At two weeks after infection, we observed an increase in *CXCR4* gene disruption in T cells collected from the CXCR4-CRISPR mice, suggesting the enrichment of CXCR4-negative cells by the selective pressure of X4-tropic HIV-1 infection (**Figure 4D**). Notably, at the same time point, the mice engrafted with *CXCR4* knockout cells exhibited ∼30-fold lower levels of plasma viremia than in the mock-treated mice (**Figure 4E)**.

**Figure 4.**
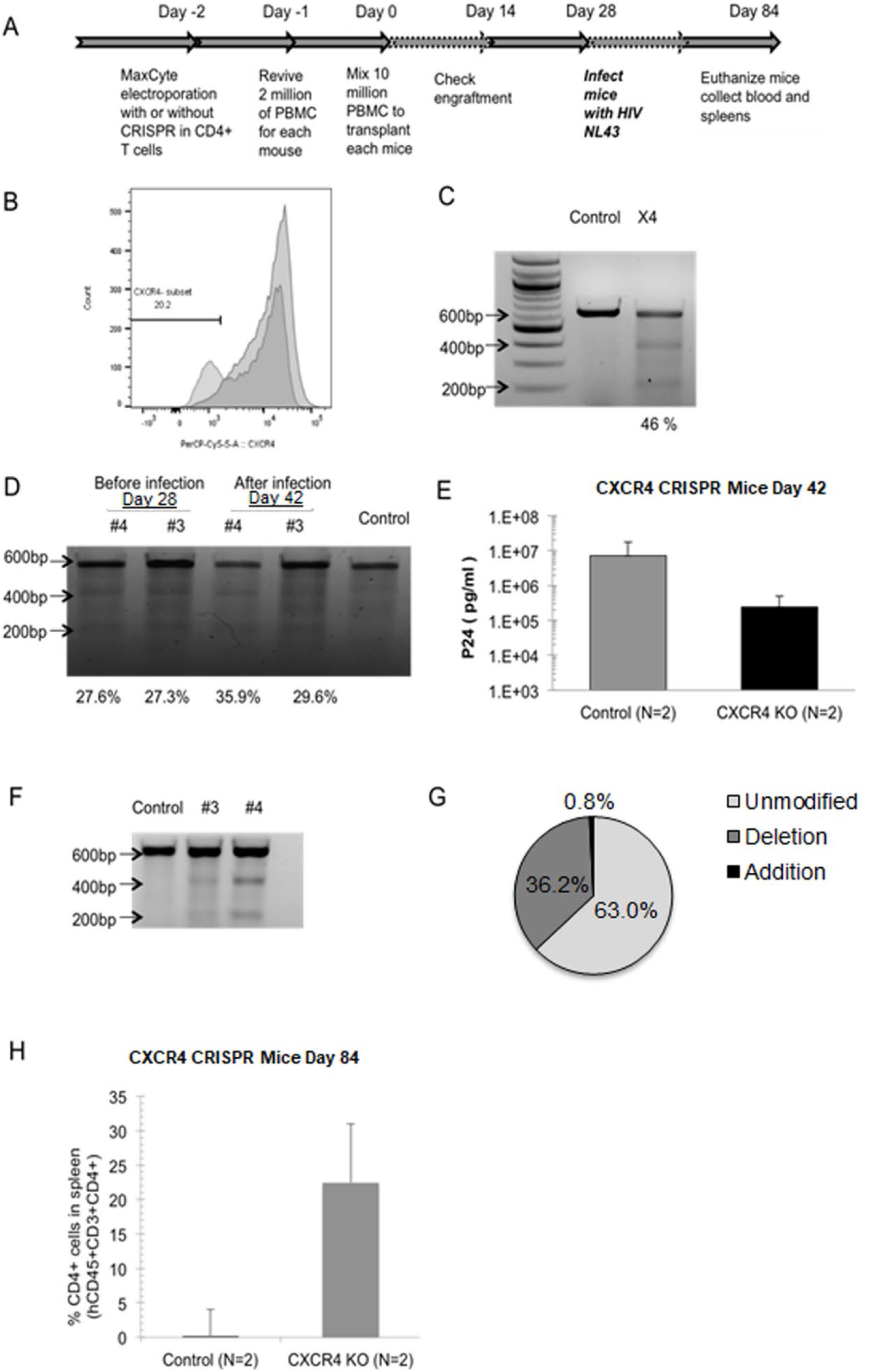
Positive selection for CXCR4 knockout cells by HIV-1_NL4-3_ infection in hu-PMBC. **A**. Schematic of the timeline of building hu-PBMC mice model and HIV infection by using mixed human primary PBMC with CXCR4 CRISPR modified CD4^+^ T cells. **B**. Cell surface CXCR4 co-receptor knockout in CD4^+^ T cells after MaxCyte electroporation of CXCR4 guide RNAs and Cas9 RNPs. Cells were fixed in 4% formaldehyde and analyzed by flow cytometry 48 hours after transfection. **C**. Surveyor assay detection of the allelic disruption of *CXCR4* gene in the CXCR4 CRISPR modified cells. **D**. Surveyor assay detection of the allelic disruption of *cxcr4* gene in the PBMC from CXCR4 CRISPR modified cells transplanted mice. Mice whole blood were collected by retro-orbital bleeding before HIV-1_NL4-3_ infection (4 weeks after transplantation) and 2 weeks after HIV-1_NL4-3_ infection (6 weeks after transplantation). **E**. qPCR was performed using plasma from hu-PBMC mice. Mice whole blood was collected by retro-orbital bleeding 2 weeks after HIV-1_NL4-3_ infection (6 weeks after transplantation). Data were presented by comparing two groups of mice which were transplanted by using the control and CXCR4 CRISPR modified cells. **F**. Surveyor assay represent the allelic disruption of *cxcr4* gene in the spleen cells from humanized mice transplanted by using CXCR4 CRISPR modified or unmodified cells (control). **G**. Quantitative analysis of indels generated by CXCR4 CRISPR in spleen cells in humanized mice. **H**. Flow cytometry analysis of CD4^+^ T cell numbers in mice spleen 12 weeks after transplantation by using CXCR4 CRISPR modified or unmodified cells (control).

At 12 weeks after transplantation (i.e., 8 weeks after HIV-1_NL4-3_ challenge), the experiment was terminated, the CXCR4-CRISPR modified cells were collected from the spleens of humanized mice (**Figure 4F)**. We analyzed the gene modification level of CXCR4-CRISPR in the mice spleens by Sanger sequencing followed by analysis using inference of CRISPR edits (ICE), which revealed 37.0% of *CXCR4* alleles were disrupted (**Figure 4G)**. Moreover, the mice engrafted with *CXCR4* knockout cells exhibited significantly higher levels of CD4^+^ T cells in the spleen (22.5% CXCR4-CRISPR or 0.2% mock-treated) than to the mice that received mock-treated cells (**Figure 4H)**. These results indicate that CRISPR-mediated gene disruption of *CXCR4* protects CD4^+^ T cells *in vivo* from infection of X4-tropic HIV-1 and virus-induced cell death.

### CCR5 and CXCR4 genome-disrupted confers primary T cells resistant broad HIV-1 infection

Specifically, we utilized the Alt-R CRISPR-Cas9 system (Integrated DNA Technologies, Inc.), which consists of spCas9 recombinant protein complexed with a trans-activating crRNA (tracrRNA) and a chemically modified CRISPR RNA (crRNA) that is specific for either *CCR5* or *CXCR4* (referred hereafter as R5X4-CRISPR). While CRISPR-mediated disruption of *CCR5* confers resistance to R5-tropic HIV-1, and disruption of *CXCR4* confers resistance to X4-tropic HIV-1, it may be necessary to edit both surface receptors to create resistance to all HIV-1 infection. To test this hypothesis, we prepared Cas9 RNP complexes with CCR5 and CXCR4 gRNAs (referred hereafter as R5X4-CRISPR) following manufacturer’s instructions. After transfection of the R5X4-CRISPR system into primary CD4^+^ T cells, we first analyzed the knockout efficacy of CCR5 and CXCR4 receptors on the cell surface. Analysis by flow cytometry revealed that the gene-modified cells exhibited a decrease in CCR5 surface expression from 88.7% in control cells to 54.9% (**Figure 5A**) and from 77.1% to 26.3% in CXCR4 expression (**Figure 5B**). In total, the proportion of dual-positive CCR5^+^CXCR4^+^ cells decreased from 85.2% to 36.8%, while levels of dual-negative CCR5^-^CXCR4^-^cells increased from 10.6% to 49.8% (**Figure S2**). This demonstrates that transient delivery of CRISPR/Cas9 is effective in knocking out both of the co-receptors that are required for HIV infection in human primary CD4^+^ T cells.

**Figure 5:**
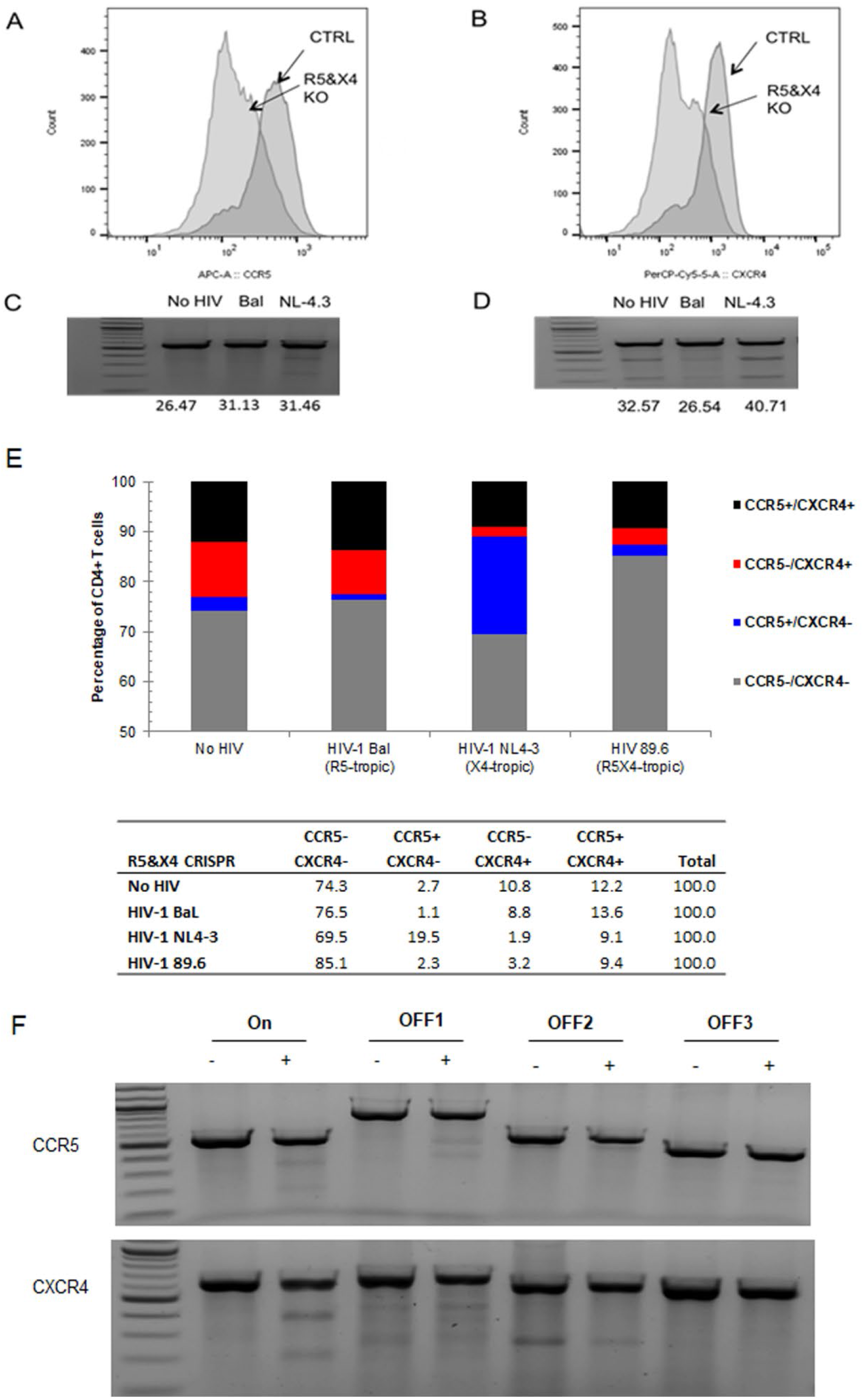
Gene disruption of CCR5 and CXCR4 leading to HIV resistance in primary CD4^+^ T cells using MaxCyte electroporation. **A-B**. Human primary CD4^+^ T cells were transfected with CCR5 and CXCR4 guide RNAs with Cas9 RNP by MaxCyte electroporation using the ‘P4’ setting. Elimination of cell surface expression of CCR5 (A) and CXCR4 (B) co-receptors are evaluated by flow cytometer. **C-D**. Surveyor assay tested CCR5 (C) and CXCR4 (D) allelic disruption in CCR5 and CXCR4 CRISPR treated cells are infected by HIV-1 virus (BaL or NL4-3). Cells were infected by using each HIV-1 virus (BaL or NL4-3) each strain after CD3/CD28 activation. 5 weeks after infection, cells were collected for genomic DNA extraction and Surveyor assay assessment. **E**. Flow cytometry analysis of CCR5 and CXCR4 expression on surface of CD4^+^ T cells treated with CCR5 and CXCR4 CRISPR and then infected by using each HIV-1 virus strain (BaL-1, NL4-3 or 89.6) after CD3CD28 activation. Cells were collected 5 weeks after infection. The bar graph represents that in the CCR5 and CXCR4 CRISPR treated cells, percentage of CCR5^-^CXCR4^-^ cells (gray bar), CCR5^+^CXCR4^-^ (blue bar), CCR5^-^ CXCR4^+^ (red bar) and CCR5^+^/CXCR4+ cells (black bar) were compared after difference strain infection. The table graph represents that in the CCR5 and CXCR4 CRISPR treated cells, percentage of CCR5^-^ CXCR4^-^ cells, CXCR4^+^CCR5^-^ cells, CXCR4^-^CCR5^+^ cells, and CXCR4^+^CCR5^+^ cells were compared after difference strain infection. **F**. Off target sites predicted by Cas-OFFinder. Top three off target gene were analyzed by Surveyor assay in the CCR5 and CXCR4 CRISPR treated cells.

We next sought to determine whether CD4^+^ T cells with disrupted *CCR5* and *CXCR4* alleles would become resistant to HIV-1 infection. We challenged the R5X4-CRISPR-modified primary CD4^+^ T cells with HIV-1 virus that utilized the *CCR5* co-receptor (HIV-1_BaL_), the *CXCR4* co-receptor (HIV-1_NL4-3_), or either the *CCR5* or *CXCR4* co-receptors (HIV-1_89.6_). Analysis of indels by Surveyor assay revealed that slight increase in disruption of the *CCR5* allele was observed after challenge with R5-tropic HIV-1_BaL_ **(Figure 5C)**. Similarly, gene disruption of the *CXCR4* allele was increased after infection with X4-tropic HIV-1_NL4-3_ (**Figure 5D**). Interestingly, cells that had surfaced expression of *CCR5* but not of *CXCR4* (CCR5^+^CXCR4-) were enriched after challenge with HIV-1_NL4-3_ (19.5%), but not after challenge with the other two strains that can utilize the CCR5 coreceptor (1.1% for HIV-1_BaL_ and 2.3% for HIV-1_89.6_). Likewise, cells with surface expression of *CXCR4* but not of *CCR5* (CCR5-CXCR4^+^) were enriched after challenge with HIV-1_BaL_ (8.8%), but not after challenge with strains that may infect via the CXCR4 co-receptor (1.9% for HIV-1_NL4-3_ and 3.2% for HIV-1_89.6_). Most notably, the CCR5^-^ CXCR4-dual-negative subpopulation increased from 74.3% in the R5X4-CRISPR cells before HIV-1 challenge to 85.1% in the cells challenged with HIV-1_89.6_, demonstrating an enrichment of cells that lack both CCR5 and CXCR4 co-receptors after incubation with this dual-tropic HIV-1 strain (**Figure 5E**).

To ascertain possible off-target gene disruption after R5X4-CRISPR treatment, we examined three possible off-target sites for the *CCR5* sgRNA target sequence and three more for the *CXCR4* target sequence, as predicted by Cas-OFFinder. Each site was analyzed using Surveyor assay, but no increases in gene disruption were observed for any of the six predicted off-target sites, whereas clear gene disruption was observed for each of the two on-target sites (**Figure 5F, Table S1, S2**).

### Poor engraftment of R5X4-CRISPR knockout CD4^+^ T cells in lymphoid tissues in Hu-PBMC mice

As shown in Figure 5, knockout of both CCR5 and CXCR4 co-receptors is necessary to block infection from R5 and X4-tropic HIV-1 strains. Thus, we tested this the R5X4-CRISPR approach in hu-PBMC NSG mice with CRISPR-induced disruption of both *CCR5* and *CXCR4*. (**Figure 6A**) After transfection of the R5X4-CRISPR RNP complex into primary CD4^+^ T cells by using MaxCyte electroporation system, we first analyzed the knockout efficacy of CCR5 and CXCR4 receptors on the cell surface. The proportion of dual-negative CCR5-CXCR4-cells increased from 21.8% to 49.0% (**Figure 6B**). Editing of the CCR5 and CXCR4 alleles was also confirmed by Surveyor assay, which revealed 22.03% and 32.03% gene disruption (**Figure 6C**).

**Figure 6.**
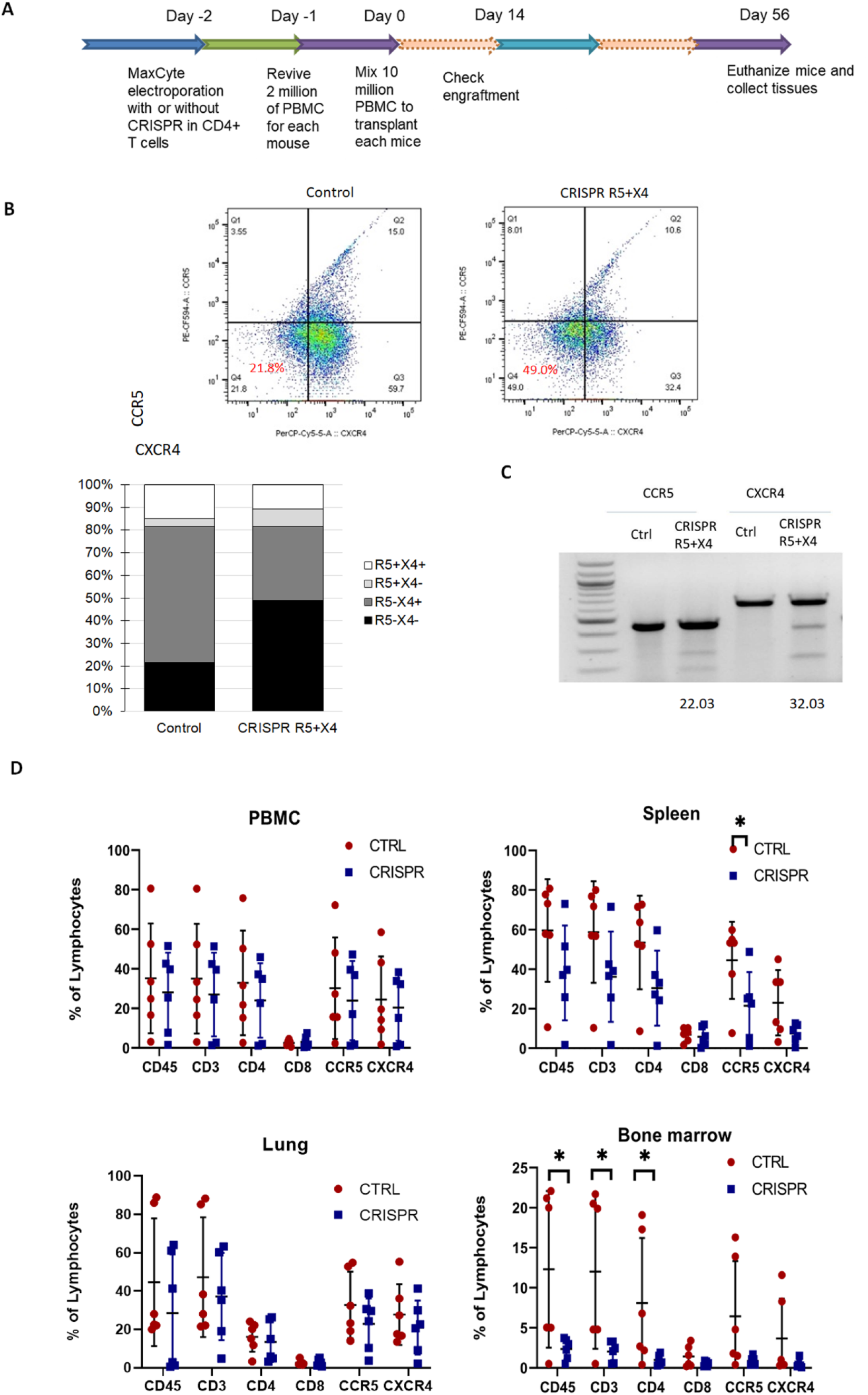
Bio-distribution of CCR5- and CXCR4-CRISPR knockout CD4^+^ T cells in Hu-PBMC mice tissues. **A**. Schematic of the timeline of building hu-PBMC mice model by using mixed human primary PBMC with CXCR4 CRISPR modified CD4^+^ T cells. **B**. Surveyor assay detection of CCR5 and CXCR4 allelic disruption in CD4^+^ T cells after MaxCyte electroporation of CCR5 and CXCR4 guide RNAs and Cas9 RNPs. **C**. Cell surface CCR5 and CXCR4 co-receptor knockout in CD4+ T cells after MaxCyte electroporation of CCR5 and CXCR4 guide RNAs and Cas9 RNPs. Cells were fixed in 4% formaldehyde and analyzed by flow cytometry 48 hours after transfection. **D**. Eight million of CRISPR modified or un-modified CD4+ T cells with 2 million human PBMCs were transplanted into NSG mice. At final time point whole PBMCs, spleens, lungs, and bone marrow of all the mice from each group were harvested and cells were analyzed by flow cytometer. (n=6, *p < 0.05)

First, we analyzed the engraftment of the gene-modified cells. From analysis of peripheral blood in the hu-PBMC mice, there were similar levels of human CD45^+^ lymphocytes or other surface markers, including CD3, CD4, CD8, CXCR4, and CCR5 between the dual CRISPR and control mouse groups (**Figure 6D**). However, we also evaluated engraftment in primary lymphoid tissues and lung to assess the homing and persistence of the CRISPR-modified cells. In the spleen, we observed slightly lower levels of CD45^+^ human cells, CD45^+^CD3^+^ T cells, CD45^+^CD3^+^CD4^+^ T cells, CD45^+^CD3^+^CD4^+^CCR5^+^ T cells, and CD45^+^CD3^+^CD4^+^CXCR4^+^ T cells in the R5X4-CRISPR-treated mice than in controls, although none of these differences was statistically significant (**Figure 6D**). Similar trends were also observed in the lung, although statistical significance was not met. However, in the bone marrow, the R5X4 mice had statistically significant (p<0.05) lower levels of human CD45^+^ cells and CD45^+^CD3^+^ T cells, as well as similar trends of slightly lower levels of CD4^+^, CCR5^+^, and CXCR4^+^ T cells (**Figure 6D**). These results suggest that CRISPR-mediated knockout of *CCR5* and *CXCR4* may alter the homing, persistence, and expansion of these cells into the bone marrow and potentially other lymphoid tissues after transplantation.

## Discussion

Owing to their essential roles as co-receptors for HIV entry and infection, the human CCR5 and CXCR4 chemokine receptors are attractive targets for gene disruption for creating HIV resistance. In this study, we investigated the versatility of the CRISPR-Cas9 in simultaneously editing both CCR5 and CXCR4 receptors human cells. We successfully disrupted *CCR5* in CD4^+^ T cell lines (**Figure 1**), primary CD4^+^ T cells (**Figure 5**), and CD34^+^ HSPCs differentiated macrophages (**Figure 3**), which all led to R5 tropic HIV-1 virus resistance. Likewise, by disrupting *CXCR4* in a CD4^+^ T cell line (**Figure 3**), primary CD4^+^ T cells (**Figure 5**), and in transplanted CD4^+^ T cells in a humanized mouse model (**Figure 4**), we achieved X4 tropic HIV-1 virus resistance.

To generate CRISPR-modified CD4^+^CCR5^-^CXCR4-T cells, we utilized the MaxCyte electroporation system, which is a scalable system that has been used for clinical manufacturing of gene-modified cells ^39^. Upon treatment with the Cas9 RNP complexes with *CCR5* and *CXCR4* gRNAs, we observed efficient gene editing for both receptors in primary CD4^+^ T cells, resulting in approximately 50% CCR5-CXCR4-double-negative cells (**Figure S3**). The gene-modified cells were resistant to broad HIV-1 infection and were selectively enriched by the selective pressure of HIV-1 infection (**Figure 4E**). In the hu-PBMC NSG mouse model, the CRISPR-modified cells were well tolerated, as the percentage of gene modified cells did not decrease over time in mice (**Figures 4D and 6D**). Moreover, in CXCR4-CRISPR humanized mice, X4-tropic HIV-1 resistance resulted in the selective enrichment of CD4^+^ T cells in spleen tissue compared to non-CRISPR mice (**Figure 4H**).

While CRISPR-mediated disruption of *CXCR4* was successful in reducing viremia and protecting CD4^+^ T cells *in vivo* (**Figure 4**), we observed that levels of R5X4-CRISPR-modified CD4^+^ T cells were significantly lower than unmodified controls in the bone marrow. CXCR4 is known to function as a surface receptor for cell homing, such as for the homing of hematopoietic stem and progenitor cells (HSPC) in the bone marrow,^40^ while the CXCR4 antagonist AMD3100 (plerixafor) is used clinically to mobilize CD34^+^ HSPCs from the bone marrow into the peripheral blood.^41^ However, it is unknown whether gene disruption of *CXCR4* would abate engraftment of CD4^+^ T cells in lymphoid organs. Previous studies have used zinc-finger nucleases to disrupt *CXCR4*^25,26^ or both *CCR5* and *CXCR4*^29^ in CD4^+^ T cells to create X4-tropic HIV-1 resistance in tissue culture and *in vivo*. Similar to our observations in Figure 4, these studies also showed decreases in HIV-1 plasma viremia and protection of the modified CD4^+^ T cells in hu-PBMC mouse models. However, these studies only evaluated CD4^+^ T cells and viremia in the peripheral blood and spleen, with no analyses of the engraftment in the bone marrow or lung. The potential toxicity of disrupting *CXCR4* in HSPCs is well established^42^, but this possibility is not necessarily associated with CD4^+^ T cells.

While gene disruption of *CCR5* continues to be evaluated clinically with promising results^32^, gene editing strategies for *CXCR4* have not advanced to the clinic. Moreover, unlike the naturally occurring *CCR5-Δ32* homozygous mutation, homozygous *CXCR4* knockouts are embryonic lethal in a murine model.^43^ Based on our observations of reduced engraftment of T cells in bone marrow following CRISPR-mediated disruption of *CCR5* and *CXCR4*, it is not clear that this strategy would be viable in humans.

## Materials and Methods

### Cell lines and viruses

CEM.NK^R^ CCR5+ cells (abbreviated as CEM-CCR5) and Jurkat cells are CD4^+^ T lymphoblastic cell lines obtained from NIH AIDS Reagent Program (catalog #4376), which is cultured in RPMI 1640 media supplemented with 10% fetal bovine serum and 2 mM L-glutamine. Human embryonic kidney (HEK) 293T cells were from ATCC (catalog #CRL-3216). HIV-1 infectious virus (HIV-1_BaL_, catalog #510; HIV-1_89.6_, catalog #1966) and molecular clone plasmid (HIV-1_NL4-3_, catalog #114), were obtained from NIH AIDS Reagent Program.

### PBMCs and primary CD4^+^ T cells

Human peripheral blood mononuclear cells (PBMCs) were isolated from leukocyte reduction system chambers (i.e., buffy cones), which were obtained from healthy human donors at the City of Hope Amini Apheresis Center (Duarte, CA). PBMCs were separated from the by centrifugation with Ficoll-Paque Premium (BD). Primary human CD4^+^ T cells were further purified and enriched by the CD4^+^ T cell isolation Kit (Miltenyi Biotech) according to the manufacturer’s instructions and then maintained in complete RPMI medium supplemented with 10% FBS.

### Guide RNA design and CRISPR-Cas9 lentiviral vector constructs

Guide RNA sequences for the *ccr5* and *cxcr4* target sites were designed using the computational tool originally described by Hsu, et al.^35^ The pL-CRISPR-SFFV-tRFP plasmid was obtained from Addgene (Plasmid #57826) and originally deposited by the Ebert lab.^34^

### Lentiviral vector production

Lentiviral vectors were packaged in HEK 293T cells by calcium phosphate precipitation. Briefly, 15 µg of transfer plasmid was cotransfected with helper plasmids (15 µg of pCMV-Pol/Gag, 5 µg of pCMV-Rev, and 5 µg of pCMV-VSVG) into HEK 293T cells with 90–95% confluency per 10-cm dish. Viral supernatant was harvested 48 hours post-transfection, concentrated by ultracentrifugation, and stored at -80°C until use. Viral titers were determined by transduction of HT1080 cells and analyzed for EGFP expression with fluorescence-activated cell sorting analysis.

### Flow cytometry analysis

To analyze cell surface expression of CCR5 and CXCR4, cells were incubated with an APC-conjugated mouse anti-human CCR5 (Becton Dickinson), PerCP-Cy5-conjugated mouse anti-human CXCR4 (Becton Dickinson) for 30 min at 4 °C. Cells then were washed twice with FACS buffer (PBS containing 1% BSA and 0.02% NaN3) and then washed twice with FACS buffer and fixed with 2% formaldehyde. FACS analysis was performed on Fortessa (Becton Dickinson, Mountain View, CA).

To isolate Tag-RFP cell populations from total CEM-CCR5 cells transduced with lentiviral vectors expressing Cas9 NLS and sgRNAs, cells were sorted using an Aria SORP cell sorter (Becton Dickinson).

### Surveyor nuclease assay

To detect indels generated by CRISPR, genomic DNA from the CRISPR modified or unmodified cells was extracted using QiAmp DNA mini Kit (Qiaqen) and assayed by Surveyor nuclease assay (Transgenomic).

### HIV-1 in vitro challenge assay

To test whether CCR5 and CXCR4 gene-disrupted cells were resistant to HIV-1 infection, cells were infected with R4-tropic HIV-1_NL-4.3_, R5-tropic HIV-1_BaL_, or dual-tropic HIV-1_89.6_ at the MOI between 0.01-0.1 at 37°C, 5% CO2 for overnight. Cells were then washed twice with PBS and re-suspended in fresh complete medium. After the challenge, cells and culture supernatants were collected every 3 days and replenished with fresh medium for a total of 28 days. Levels of HIV-1 gag p24 in culture supernatants were measured by ELISA as instructed by manufacturer (PerkinElmer).

### Generation of adult HSPC-derived macrophages

Cord blood was purchase from StemCyte (Baldwin Park, CA) with approval from the City of Hope Institutional Review Board (IRB 17155). Sorted CD34^+^ HSPCs were cultured in Iscove’s modified Dulbeco’s media with 20% FBS supplemented with 2 mmol/l of glutamine, 25 ng/ml of stem cell factor (Stemcell Tech), 30 ng/ ml of Flt3-L (PeptroTech), 30 ng/ml of interleukin-3 (Gibco), and 30 ng/ml of macrophage colony stimulating factor (PeproTech, Rocky Hill, NJ) for 10 days for guided differentiation to monocytes and were then switched to DMEM with 10% FBS supplemented with 2 mmol/l of glutamine, 10 ng/ml of granulocyte macrophage colony stimulating factor (PeproTech), and 10 ng/ml of macrophage colony stimulating factor (PeptroTech) for 5 days for activation into macrophages. Adherent macrophage cells were collected for HIV challenge experiments. The purity of cells was typically greater than 90% CD14^+^ based on fluorescence-activated cell sorting analysis.

### Primary CD4^+^ T cell electroporation

The transfection of primary CD4^+^ T cells was performed on MaxCyte STX. 2×10^7^ primary CD4^+^ T cells were centrifuged and washed twice with 1x PBS, and the cell were re-suspended with 100 μl prepared EP buffer and Cas9 NLS and chemically modified guide RNA with tracrRNA complex ordered form IDT. The mixture was then transferred to the OC-100 cuvette and electro-transfected with MaxCyte STX programs. After transfection, the cells were transferred to a CD3/ CD28 coated six well plate and cultured with RPMI 1640 supplemented with 10% FBS, and IL-2 (100 IU/ml).

### Humanized PBMC (hu-PBMC) NSG mouse model

NOD.Cg-*Prkdc*scid IL2rgtm1Wjl/SzJ (NSG) mice were obtained from The Jackson Laboratory (Bar Harbor, ME) and bred at the City of Hope Animal Resources Center according to the protocols approved by the Institutional Animal Care and Use Committee of the City of Hope (IACUC 16095). Adult NSG mice at age of 8–10 weeks old were transplanted with human PBMCs via intraperitoneal injection. Specifically, each mouse received 2.0 × 10^6^ human PBMCs mixed with 8.0 × 10^6^ CRISPR modified or un-modified human CD4+ T cells.

### HIV-1 qRT-PCR

HIV-1 viral RNA was extracted from 20-50 ul of plasma using QIAamp Viral RNA mini kit (Qiagen). qRT-PCR was performed using a TaqMan Fast Virus 1-Step Master Mix, according to the manufacturer’s instructions (Applied Biosystems, Foster City, CA). The primers used were LTR-F (5’-GCCTCAATAAAGCTTGCCTTGA-3’) and LTR-R (5’-GGCGCCACTGCTAGAGATTTT-3’), along with a probe (5’-FAM/AAGTAGTGTGTGCCCGTCTGTTGTGTGACT-3’). Assay was performed using automated CFX96 TouchTM Real Time PCR Detection System (Bio-Rad).

### Off-target analysis

Cas-OFFinder was employed to find potential OTSs with limitation of three-base mismatched sequences. From the resulting off-targets, OTSs only in gene-coding regions were selected and Surveyor nuclease assayed (Surveyor Mutation Detection Kit; Transgenomics).

### Deep sequencing and CRISPResso analysis

Target loci were amplified by the specific primers. Before sequencing on an Illumina HiSeq 2500 platform, the amplicons were purified, end-repaired and connected with sequencing primer. For the sequences gained by sequencing, low quality and joint pollution data were removed to obtain reliable target sequences (clean reads) for subsequent analysis. The corresponding Read1 and Read2 (sequences gained from the 5’- and 3’-ends, respectively) were spliced. Analysis of indels was performed using the CRISPResso tool ^44^.

## Acknowledgements

This research was supported by grants from the California HIV/AIDS Research Program (CHRP) IDEA (Innovative Developmental Exploratory Award) in Basic Biomedical Sciences (ID13-BRI-540) to J.C.B, City of Hope Analytic Cytometry Core, City of Hope Animal Resource Center. The following reagents were obtained through the NIH AIDS Reagent Program, Division of AIDS, NIAID, NIH: Jurkat cells, CEM.NK^R^ CCR5+ cells from Dr. Alexandra Trkola^36^; HIV-1_Ba-L_ from Dr. Suzanne Gartner, Dr. Mikulas Popovic and Dr. Robert Gallo^45^; HIV-1_89.6_ Virus from Dr. Ronald Collman^46^; and HIV-1_NL4-3_ Infectious Molecular Clone (pNL4-3) from Dr. Malcolm Martin^47^. Cord blood from anonymous donors was purchased from StemCyte.

## Authors’ contributions

JB, SL: Conceived and designed the experiments.SL: Performed the experiments. LH: Helped with mouse tissue collection. JB, SL: Analyzed the data. SL, JB: Wrote the paper. All authors read and approved the final manuscript.

## Notes

### Competing Interest Statement

The authors have declared no competing interest.

