## Supplemental Files for "CRISPR/Cas9-mediated gene disruption of endogenous co-receptors confers broad resistance to HIV-1 in human primary cells and humanized mice"

### **Supplemental Material**

Supplemental Figure 1- Deep sequencing analysis of CCR5 CRISPR generated indels in CCR5+CD4+CEM T cells.

Supplemental Figure 2- Double knockout population in CRISPR modified CD4+ T cells by using MaxCyte electroporation.

Supplemental Figure 3- ICE analysis of indels generated by CRISPR CXCR4 modification in mice spleen cells.

Supplemental Table 1- CCR5 target sequences, off target sequences and PCR primer sequences used for Surveyor assay after MaxCyte electroporation

Supplemental Table 2- CXCR4 target sequences, off target sequences and PCR primer sequences used for Surveyor assay after MaxCyte electroporation

**Supplemental Figure1. Deep sequencing analysis of CCR5 CRISPR generated indels in CCR5+CD4+CEM T cells.** After CCR5 CRISPR transduction, cells were sorted by gating of Tag-RFP. Genomic DNA was extracted and primers were designed to amplify about 150bp amplicons with the cutting site in the middle. PCR product was purified and sent for next generation Sanger sequencing. Yellow highlights the PAM sites. Indels have more than 1000 reads are listed. A. Sequences of insertions sequences with more than 1000 reads. B. Sequences of deletions and/or substitutions with more than 1000 reads.

**A.**

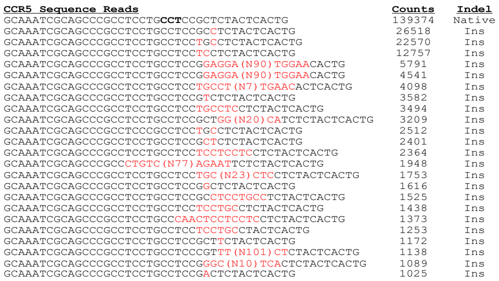


**B.**

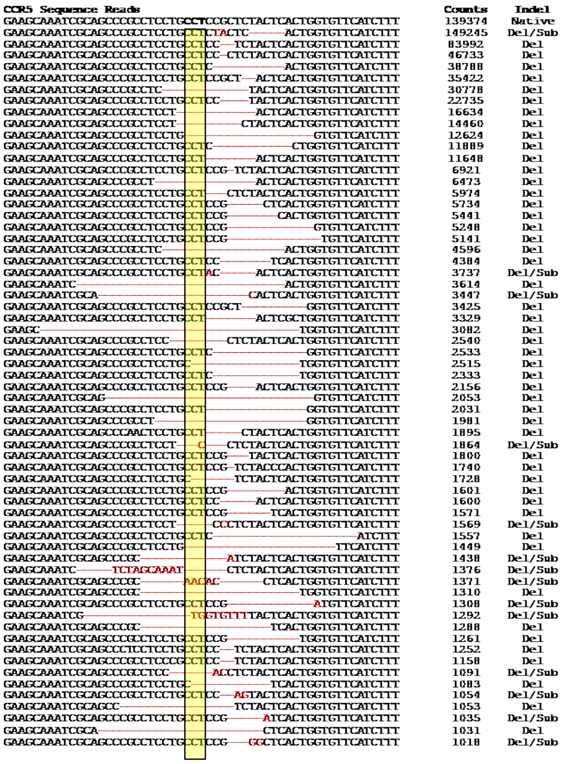

**Supplemental Figure2. Double knockout population in CRISPR modified CD4+ T cells by using MaxCyte electroporation.** In the flow cytometry data matrix, Q1 represents CCR5+CXCR4-cells, Q2 represents CCR5+CXCR4+ cells, Q3 represents CCR5-CXCR4- cells, Q4 represents CCR5-CXCR4-cells. MaxCyteGXT program 2, 4 and 6 were tested for optimal transfection efficiency.

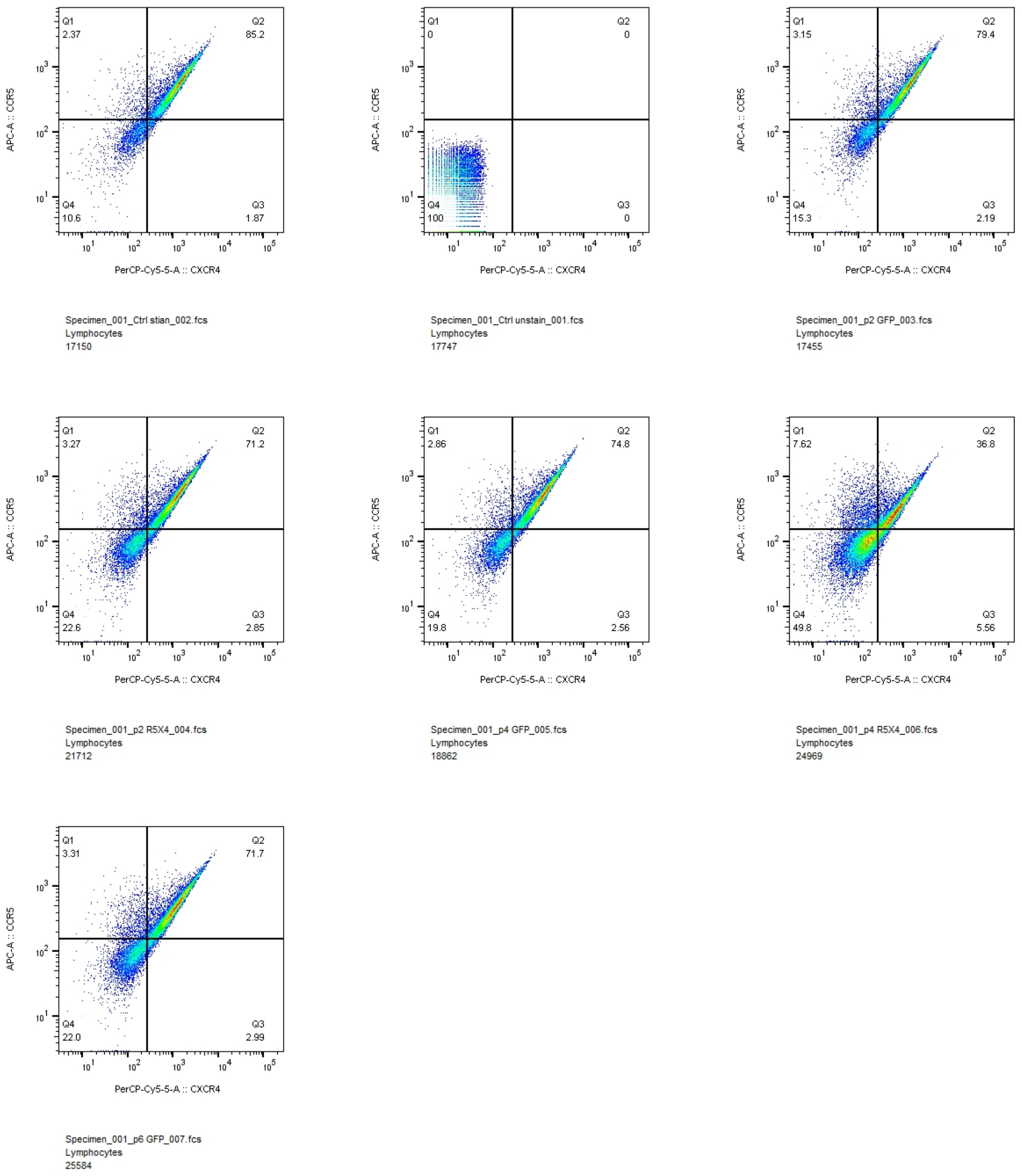

**Supplemental Figure3. ICE analysis of indels generated by CRISPR CXCR4 modification in mice spleen cells.** Mice spleens are harvested 12 weeks after transplantation which is 8 weeks after HIV-1 infection. Genomic DNA were extracted by using QiaAmp mini DNA kit and the sequences were PCR amplified by Phusion High-fidelity polymerase. PCR products are purified by using Qiagen PCR purification kit and then Sanger sequenced. The Sanger sequencing results are uploaded into SyntheGo ICE Sanger sequencing analysis system for analysis.

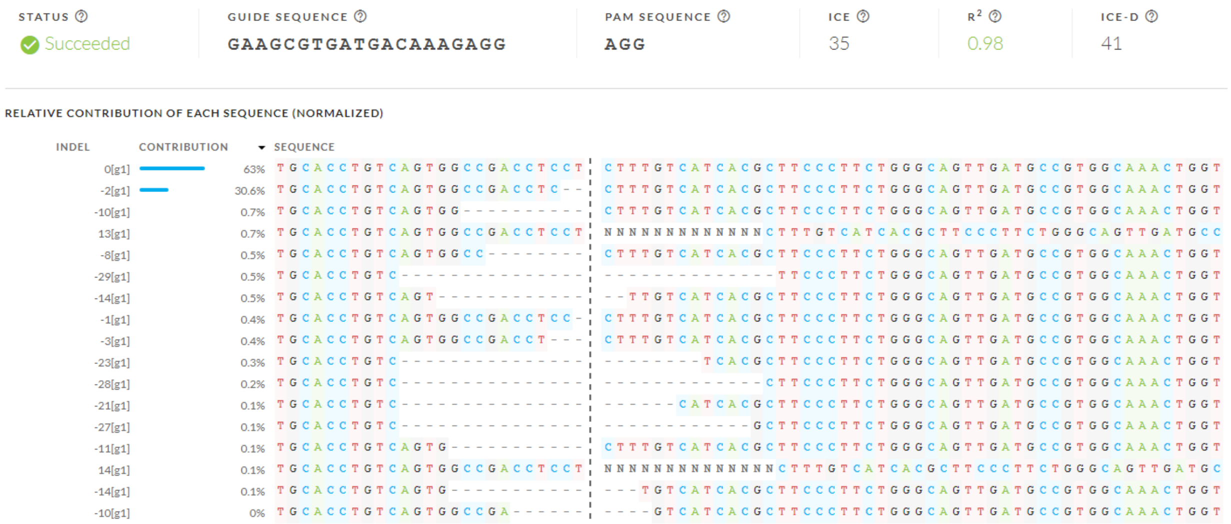

**Supplemental Table1. CCR5 target sequences, off target sequences and PCR primer sequences used for Surveyor assay after MaxCyte electroporation**

| Target name | Target sequence | Chromosome | PCR primers 5'-3' |
|---|---|---|---|
| R5 | **AACACCAGTGAGTAGAGCGGAGG** | chr3 | Forward: agcacaagattttatttggt<br>Reverse: aatagagccctgtcaagagt |
| OFF1 | **AACACCAGcGAGTAGAGCGGAGG** | chr3 | Forward: ggttctcttgtgtctgtctta<br>Reverse: tccatcctcgtggaaaataag |
| OFF2 | **AACACCAGgGAGTAcAGCGGGGG** | chr3 | Forward: atctgtatccccattcttcacca<br>Reverse: tcgattgtcagcaggattatga |
| OFF3 | **AACtCCAGTGAGgAGAGgGGTGG** | chr3 | Forward: gtgaggcatcgaacagaact<br>Reverse: gatgtgcagatgcatgtcat |

**Supplemental Table 2. CXCR4 target sequences, off target sequences and PCR primer sequences used for Surveyor assay after MaxCyte electroporation**

| Target name | Target sequence | Chromosome | PCR primers 5'-3' |
|---|---|---|---|
| X4 | **GAAGCGTGATGACAAAGAGGAGG** | chr2 | Forward: cttcagataactacaccgag<br>Reverse: agcttggagatgataatgca |
| OFF1 | **GAgGCCTcATGACAAAGAGGGGG** | chr2 | Forward: tgagcactgagtatggatct<br>Reverse: agactatccatcaggacaca |

| | | | |
|---|---|---|---|
| OFF2 | **GAAGgGaGATGACAcAGAGGGGG** | chr2 | Forward: aacaggtcttgaaaggtgaa<br>Reverse: aaccaccactaacaacaaag |
| OFF3 | **GAAaCgTGATGtCAcAGAGGAGG** | chr2 | Forward: gcagggcaataaactgaact<br>Reverse: atgcgtttgcatttctgtgg |
